# Hydrogen evolution and absorption phenomena in plasma membrane of higher plants

**DOI:** 10.1101/2020.01.07.896852

**Authors:** Xin Zhang, Zhao Zhang, Zehua Su, Qinghui Zhao, Tyler W. LeBaron, Qinjian Li, Baobei Lyu, Yao Mawulikplimi Adzavon, Guangyang Liu, Mengyu Liu, Fei Xie, Pengxiang Zhao, Xuemei Ma

## Abstract

Molecular hydrogen (H_2_) exhibits a wide breadth of botanical effects especially in improving the resistance to abiotic stress. However, the primary target is not clear. Bidirectional hydrogenases have been well studied in microbes and alga, but there is a paucity of research in higher plants. Here, we used real-time spectrophotometry, gas chromatography and H_2_ sensor to detect the H_2_-evolving and H_2_-uptake activity of the plasma membrane (PM) vesicles of higher plants. Net H^+^, Na^+^ and K^+^ fluxes were measured using Non-invasive Micro-test Technology (NMT). We found that the PM vesicles of *Vigna radiata* hypocotyls and *Capsicum annuum* stems have the ability of both oxidizing and producing H_2_. This two-way enzymatic activity coupled with the redox of ferricyanide/ferrocyanide and NADH/NAD^+^. H_2_ affected the H^+^ and Na^+^ fluxes on the surface of *V. radiata* hypocotyls. These results demonstrate that the PM vesicles of higher plants have the abilities of H_2_ evolution and absorption. The bidirectional activities may act as a valve to regulate the energy balance in plant cells.

## Introduction

Molecular hydrogen (H_2_) has been reported to have antioxidant-like effects by effectively protecting cells against oxidative stress [1]. Additional research has demonstrated that H_2_ exhibits a wide breadth of functions including anti-inflammatory, anti-allergy, anti-apoptosis, and anti-metabolic disorders[2]. In addition to the beneficial roles of H_2_ reported in animals, more than 30 studies have reported that it also has an important regulatory effect on the physiological function of plants. H_2_ could enhance the plant tolerance and resistance to several abiotic stresses such as high salinity, cadmium, paraquat exposure, heat stress, drought tolerance, as well as delaying of kiwifruit ripening[3]. H_2_ appears to improve redox homeostasis via modulating reactive oxygen and nitrogen species (ROS), which mediates many of the biological effects of H_2_ [4].

It’s well known that microbes ranging from ancient bacteria to eukaryotic microalgae are capable of metabolism H_2_. Their ability to either produce or use H_2_ as a metabolite arises from the expression of hydrogenase enzymes [5]. Hydrogenases are the group of diverse metalloenzymes that catalyze the reversible reaction of dihydrogen into protons and electrons: H_2_ (⇆ H^+^+ H^−^) ⇆ 2H^+^ + 2e^−^. The two main types of hydrogenases are classified by the metal clusters at their catalytic sites and are termed [FeFe] and [NiFe]-hydrogenases. However, only a few studies have reported H_2_ metabolism in higher plants.

It was hypothesized that hydrogenase enzymes exist in some higher plants in 1947 [6]. Sanadze firstly provided evidence that the leaves of higher plants could both release and absorb H_2_ [7]. Then Renwick’s group demonstrated that many kinds of higher plants could generate H_2_ [8]. In 1984, Torres V. et al. reported experimental evidence for hydrogenase activity in some Gramineae palts. The germinated seeds of barley produced considerable amounts of H_2_ after anaerobic treatment. The H_2_ metabolism was due to plant tissues and not to contaminant microorganisms [9]. Subsequent studies demonstrated H_2_ generation from higher plants during a seed’s germination or abiotic stresses in tissue lysis, e.g. *barley, Arabidopsis, rice, Medicago sativa* [10-13]. Zeng et al. confirmed the H_2_ production with the consistent H_2_-evolving hydrogenase activity in rice seedling lysis [14]. Compared with H_2_ evolution, H_2_ uptake in higher plants have been seldom reported. Except for chloroplasts [15], there were no other reports about the H_2_ metabolism sites of higher plants.

In this study, we investigated the H_2_-evolving and H_2_-uptake bidirectional activities in the plasma membrane (PM) vesicles of *Vigna radiata*. We demonstrate that a “hydrogenase-like complex” existed in PM acts as an electron/proton provider by oxidation of H_2_ or an electron acceptor by production of H_2_, which may help maintain the plant in redox homeostasis. Therefore, taking all the reports of physiological effects of H_2_ in higher plants, coupled with our novel discoveries, we propose that the variety of functions of H_2_ may mediated by a bidirectional H_2_ metabolism activity.

## Results

### Detection of hydrogenase activities in PM vesicles of higher plants

To investigate the H_2_-evolving ability of the PM vesicles in higher plants, crude PM vesicles of *V. radiata* hypocotyls were used firstly. Endogenous H_2_ was detected by the method of headspace GC. As shown in Figure 1A, the crude PM could release about 100 µmol/L/h/mg protein under normoxic condition and hypoxia could significantly promoted the H_2_ production for about 146 µmol/L/h/mg protein. Additionally, we found that metal ion chelators EDTA can completely inhibit H_2_ generation (Figure 1B), which indicated that the enzyme belongs to the class of metalloenzyme. Then the H_2_ sensor electrode was used to monitored the H_2_-evolving hydrogenase activity of purified PM vesicles continuously. As indicated in figure 1C, the time-course of changes in H_2_ levels was monitored and the maximum cumulative H_2_ production was 57 µmol/L/mg protein (Figure 1C and Figure S1A), but no H_2_ evolution was observed in boiled PM vesicles (Figure S1A). Except *V. radiata*, H_2_ evolution was also observed in purified PM vesicles from *C. annuum* stems and the maximum H_2_ production activity was about 314 µmol/L/mg protein (Figure S1B). Above results indicated that the PM vesicles of higher plants were capable of producing H_2_ endogenously.

**Figure 1.**
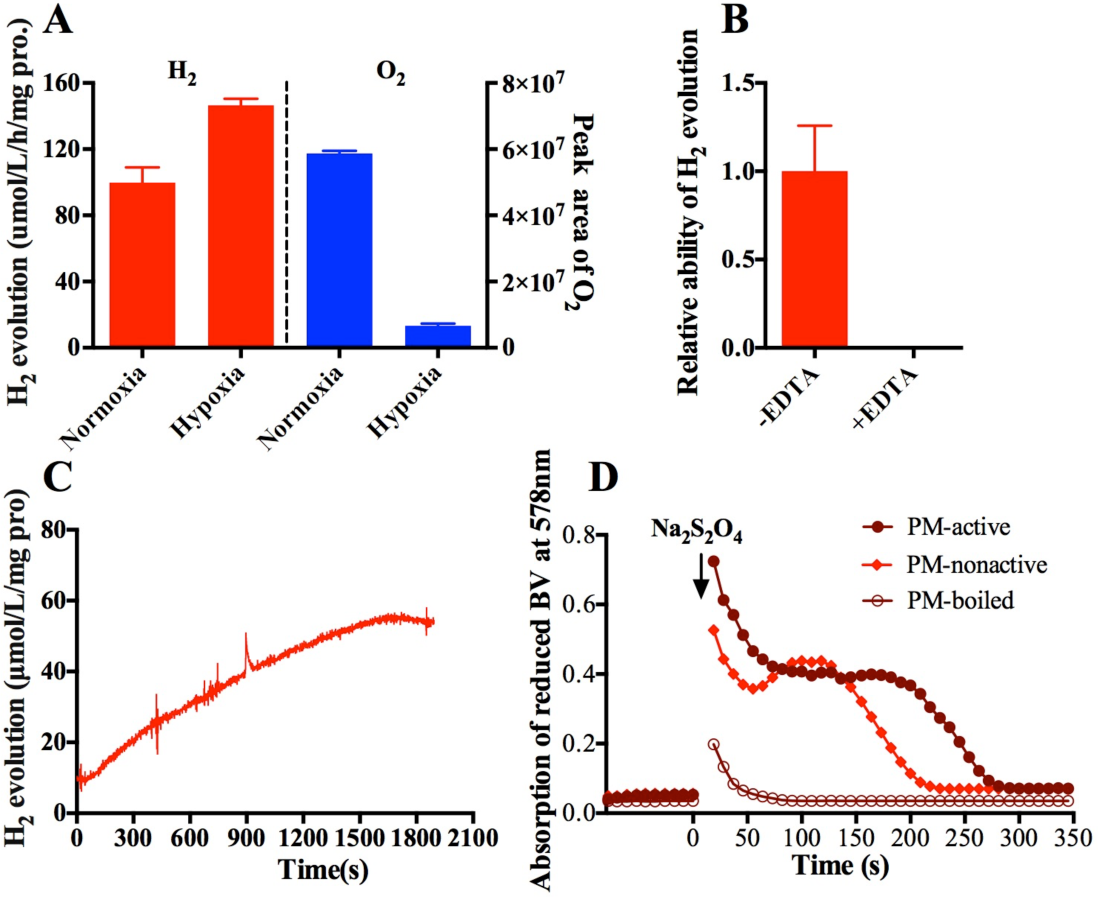
The bidirectional hydrogenase activities of PM vesicles from *V.radiata* hypocotyls. **(A)** The effect of O_2_ on H_2_ evolution of crude PM vesicles. **(B)** The effect of EDTA on H_2_ evolution of crude PM vesicles. **(C)** The H_2_ evolution activity of purified PM detected by H_2_-sensor. **(D)** The time-course of H_2_ uptake activities of purified PM vesicles. PM vesicles were either active or non-active with or without 5 µM NADH. Na_2_S_2_O_4_ initiated the reactions as indicated by arrows.

We further investigated whether purified PM vesicles were capable of utilizing H_2_ as an electron donor to reduce terminal electron acceptor. As shown in Figure 1D, the reduced BV of PM-boiled control was oxidized quickly, whereas the other groups could delay oxidation of reduced BV. The similar results were obtained in purified PM of *C. annuum* stems (Figure S2). These results indicated that H_2_ could be used to provide electrons to BV. Additionally, the enzymatic activity of H_2_-dependent BV reduction was significantly enhanced when the PM was treated with 5 µM NADH (Figure 1D and Figure S2).

### Other enzymatic characterization of hydrogenase activity in PM

To further confirm this hydrogenase-like activity, we detected the H_2_: NAD^+^ redox reaction and NAD^+^ was assayed as an electron acceptor. In our experiment, we not only observed that the PM vesicles could utilize H_2_ to regenerate NADH, but also found that NADH was oxidized again (Figure 2A, B). Hydrogen accelerated the reversible redox reaction between NAD^+^ and NADH. These bidirectional redox reactions were the most evident when the PM was activated by 5 µM NADH (Figure 2A) compared with non-activated PM (Figure 2B).

**Figure 2.**
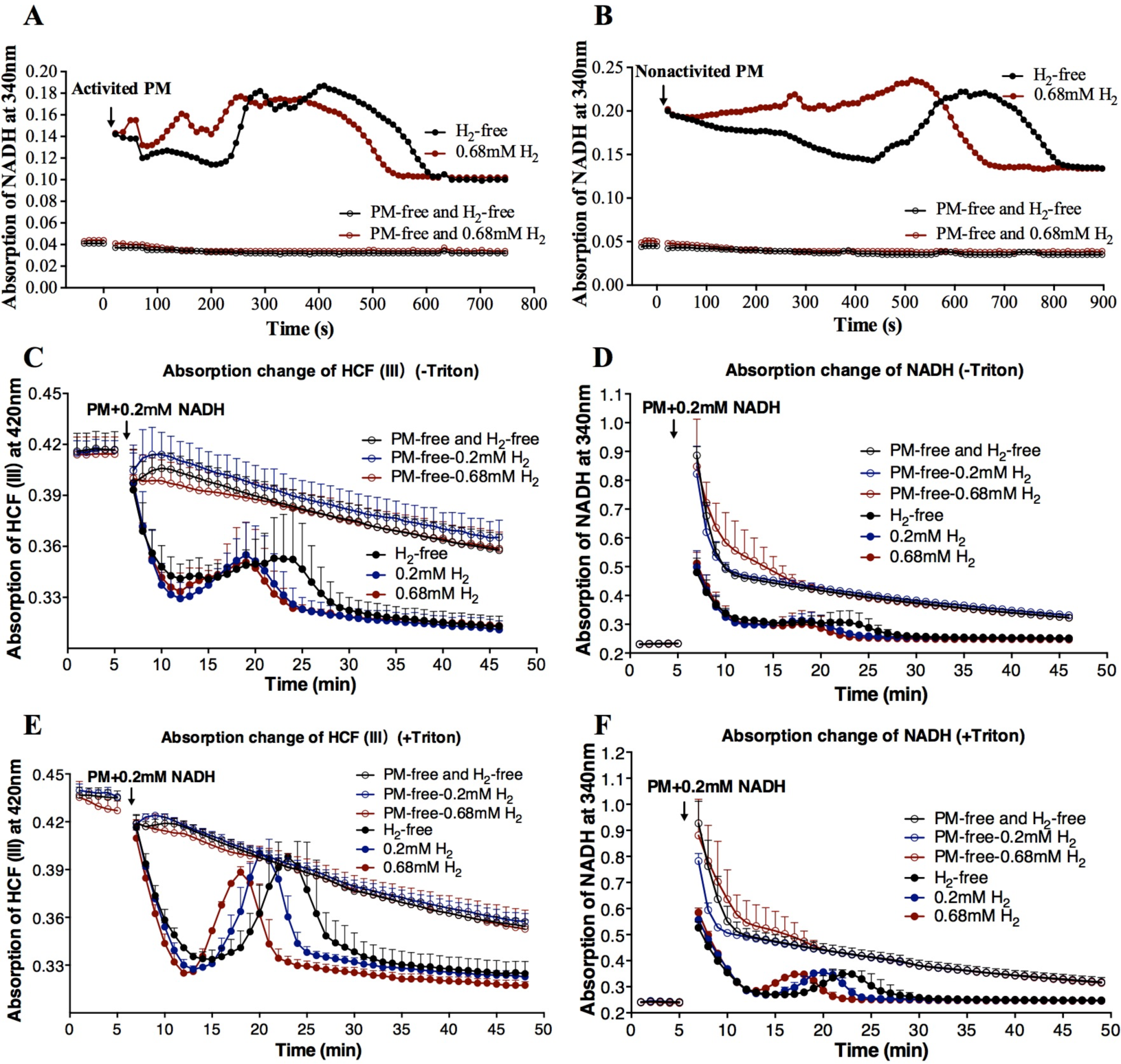
Other subunits redox activities of hydrogenase were measured by time-course spectrophotometric methods. **(A, B)** Time-course of absorption change detected for NADH (340 nm) in *V.radiata*. The PM vesicles were activated by 5 µM NADH **(A)** or not **(B)**. The reaction was initiated by addition of 10-20 µg PM. **(C to F)** Time-course of absorption change detected for HCF III (420 nm) and NADH (340 nm) simultaneously by the addition of 10-20 µg PM and 0.2 mM NADH as indicated by arrows. PM was pretreated in the absence **(C, D)** or presence **(E, F)** of 0.05% triton X-100 for 10 min to change the permeability of PM vesicles. Data shown are the means ± SD (n=3). PM-free, without PM. H_2_-free, PM was added into reaction buffer without H_2_. Black color represents that the reaction buffer did not contain H_2_. Blue and red colors represent that the reaction buffer contained 0.2 mM or 0.68 mM H_2_.

In this study, the activity of NADH-ferricyanide oxidoreductase (NFORase) of PM vesicles from *V.radiata* hypocotyls was also measured. Ferricyanide (hexacyanoferrate (III), HCF III) was used as an electron acceptor and its reduced product was HCF II, which is convenient for the investigation of electron flow[16]. The reaction of HCF III + NADH → HCF II + NAD^+^ was detected spectrophotometrically in this experiment. The process represented HCF III→HCF II→HCF III→HCF II and NADH→NAD^+^→NADH→ NAD^+^ conversion, which indicates a type of bidirectional redox reaction (HCF III + NADH ⇆ HCF II + NAD^+^) (Figure 2C to F). Triton X-100 treated permeable vesicles showed a steeper decrease and increase in absorbance (Figure 2E and F) compared with no triton treatment (Figure 2C and D). Similar to above results, the PM vesicles themselves also catalyzed the redox reaction between NAD^+^/NADH and HCF III/HCF II. H_2_ significantly accelerated the enzyme activity in a dose-dependent manner (i.e. 0.68 mM H_2_ >0.2 mM H_2_ > H_2_-free) during both the ascent and descent processes (Figure 2C to F).

### HRW influenced the H^+^ fluxes in *V. radiata*

To determine whether H_2_ had the influence on the growth of plants, the seeds of *V. radiata* were germinated in hydrogen-rich water (HRW) in darkness. H_2_ could accelerate the elongation of *V. radiata* hypocotyls significantly (Figure 3A and 3B), but not its diameter (Figure 3C). The hypocotyls elongation was due to a significant extension of the cortex cells (Figure 3D). The elongation of plant cells can be promoted by H^+^ secretion according to the acid growth hypothesis. Therefore, to confirm whether the elongation was due to the change of H^+^ fluxes on the hypocotyls surface influenced by H_2_, NMT technique was used. During growth, plants rely on the roots to absorb HRW, so real-time H^+^ fluxes of hypocotyls were measured to simulate the actual situation of using H_2_ while roots were treated with water which contained dissolved H_2_ or not (Figure 3E). As shown in Figure 3F and 3G, the net H^+^ efflux of hypocotyls was significantly enhanced by 0.8 mM H_2_.

**Figure 3.**
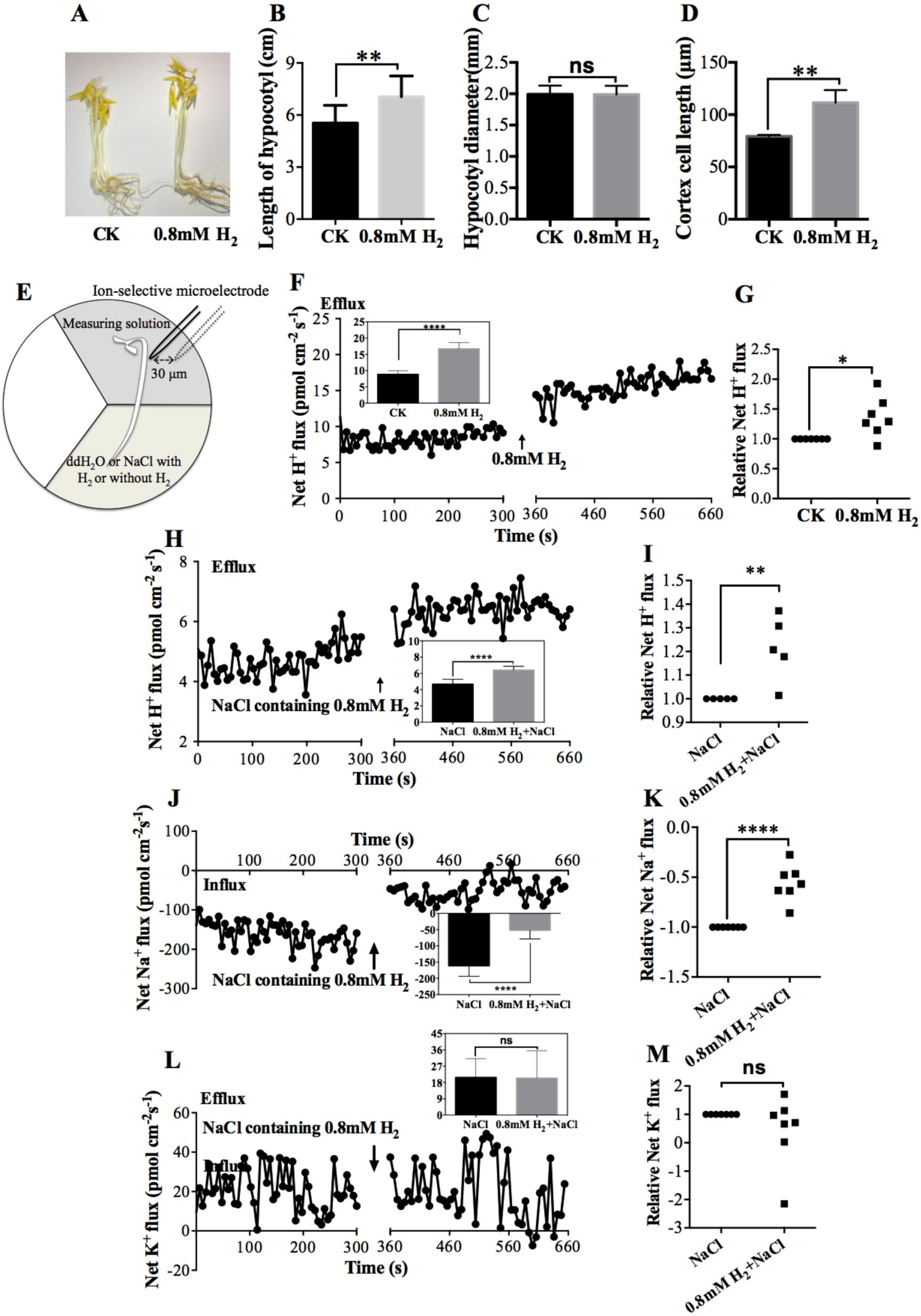
Biological effects of H_2_ on *V. radiata*. **(A-D)** Hydrogen’s biological effects on the growth of seedlings **(A)**, the length of hypocotyls (mean ±SD, n=12) **(B)**, the diameter of hypocotyls (mean ±SD, n=15) **(C)** and the cortex cell length of hypocotyls (mean ±SD, n=4) **(D)**. CK, H_2_-free water. **(E)** Schematic diagram of net H^+^-fluxes measurement of hypocotyls from intact *V. radiata* seedlings by NMT. **(F)** Net H^+^ fluxes of the hypocotyls influenced by 0.8 mM H_2_. The net H^+^ fluxes from a representative sample are shown. Steady ion fluxes without H_2_ treatment were examined for 5 min, after which the treatment solution of roots was replaced by HRW as indicated by arrows. The mean ion fluxes within the measuring periods of treatment with or without H_2_ are shown in illustrations. **(G)** Relative net ion fluxes of 7 samples are shown. **(H, J and L)** 0.8 mM H_2_ effects on net H^+^ **(H)**, Na^+^ **(J)** or K^+^ **(L)** fluxes of the hypocotyls under 150 mM NaCl were detected. The net ion fluxes from a representative sample are shown. Steady ion fluxes with NaCl treatment were examined for 5 min, after which the solution was replaced by hydrogen-rich NaCl solution as indicated by arrows. The mean ion fluxes within the measuring periods of H_2_-free or H_2_ treatment are shown in illustrations. **(I, K and M)** Relative net ion fluxes of 5-7 samples under 150 mM NaCl treatment are shown. * P<0.05,** P<0.01, **** P<0.0001.

Net H^+^ fluxes of excised hypocotyls treated directly by H_2_ were also measured (Figure S3A) and there was no significant influence by either 0.2 mM or 0.8 mM H_2_ (Figure S3B and S3C). This discrepancy compared with the intact seedlings might due to the tissues condition and the way of H_2_ treatment. The excised hypocotyls were surrounded by H_2_ and H_2_ could be used by both surface and inner of hypocotyls. If the concentrations of H^+^ inside and outside the hypocotyls changed at the same time, the change of H^+^ flow probably could not be observed.

To further investigate the H_2_ influence when plants faced abiotic stress, the H^+^, Na^+^, and K^+^ fluxes of hypocotyls under salt stress were detected. As shown in Figure 3H and 3I, H_2_ promoted H^+^ effluxes that were similar with the above results. Except for changes in H^+^ effluxes, H_2_ also significantly reduced Na^+^ influx (Figure 3J and 3K), but had no influences on K^+^ flux (Figure 3L and 3M). We suggested that the transport of salt ions across the membrane could be affected by the H_2_-uptake activitiy in PM vesicles, which could split H_2_ into protons.

## Discussion

It has been shown that exogenous H_2_ has positive botanical effects on plant growth and abiotic stress adaptability but with limited knowledge about its primary targets[3]. The PM plays an important role in plant responses to the environment in order to survive stresses since almost all environmental changes are initially perceived by the PM [17]. The investigation of H_2_ metobolism in PM vesicles could help to provide insights into the underlying molecular mechanisms responsible for the extensive biological effects of H_2_ on higher plants.

### H_2_-evolving and H_2_-uptake activities existed in PM vesicles of higher plants

All the catalytic properties detected in our study demonstrate that the PM vesicles contain reversible and functional H_2_ metabolism activities. The classical methods in detecting H_2_ uptake and H_2_ evolution by hydrogenase enzymes were used. H_2_ could be oxidized directly to provide electrons to BV, NAD^+^ or HCF (III) (Figure 1D and Figure 2). The enzymatic activity of H_2_-dependent BV reduction was significantly enhanced by 5 µM NADH (Figure 1D, Figure S2). This property was consistent with hydrogenase, which can be activated by small amounts of NADH [18]. Conversely, protons could be reduced to H_2_ by accepting electrons from reduced MV (Figure 1C, Figure S1). Previous studies [18, 19] have shown that NAD^+^-reducing hydrogenase has the module-specific enzymatic activities, e.g., the oxidation of NAD(P)H with ferricyanide as electron acceptor. In our study, the NFORase activities were found in PM vesicles of *V. radiata* and primary active sites maybe located on the cytoplasmic side of PM, which is consistent with previous studies [20]. Importantly, our study reports the bi-directional circulation of NAD^+^↔NADH and HCF (III)↔HCF (II), and H_2_ could accelerate both of these redox reactions (Figure 2). These results suggest that the module-specific enzymatic activities of NAD^+^-reducing hydrogenase also existed in PM vesicles and H_2_ participates in the bidirectional redox.

The redox state of cells, especially the level and ratio of NAD^+^/NADH, directly affects the vital processes of organism growth, aging, and death. The hydrogenase-like activities in PM vesicles is likely involved in the cellular NAD(H) pool. The hydrogenase of *Ralstonia eutropha* could couple H_2_ oxidation with the reduction of NAD^+^ to NADH, conversely it also catalyzes the formation of H_2_ from NADH to prevent ‘over-reduction’ of the cytoplasm under O_2_-limiting conditions [21]. It is conceivable that the “valve” effect of bidirectional activities of H_2_-oxidation and H_2_-evolution plays an important role in regulating the energy homeostasis of plants. Researchers have found that some abiotic stresses increase endogenous H_2_ production in higher plants, and H_2_ could alleviate damage induced by abiotic stresses [12, 13]. Wu *et al*. reported that H_2_ alleviated Cd stress through ROS detoxification by reestablishing glutathione homeostasis [22]. Chen found that HRW pretreatment remarkably attenuated heat stress-induced lipid peroxidation and antioxidant dysregulation in cucumber leaves [23]. Many botanical effects of H_2_ in plants are at least partially related to the reestablishment of redox homeostasis under abiotic stress. We speculate that the primary mechanism of how H_2_ promotes plant resistance is due to the functional activity of bidirectional H_2_-oxidation and H_2_-evolution. When energy (electron density) is high, H_2_ may be produced, which may also regulate reactive oxygen species (ROS). On the other hand, when energy is low, H_2_ can be activated by the hydrogenase to produce energy.

### H_2_ metabolism of PM can influence the transmembrane ion flow

Hydrogenases are involved in membrane-linked energy conservation and ion transportation through the generation of a transmembrane proton force [24]. In higher plants, membrane electrochemical potential gradient is closely related with a variety of biological functions such as ion transport, signal transduction, nutrient uptake, cell elongation, cellular pH regulation, etc. [25]. On the basis of homology modules between hydrogenases and complex I of the respiratory chain, hydrogenases are proposed to work as an ion (H^+^ or Na^+^) pump [26, 27]. The mechanism of energy conservation by Na^+^/H^+^ translocation has been best studied with [NiFe] hydrogenases from methanogens [28, 29].

In our research, we designed a simple but very practical method to measure net ion fluxes of hypocotyls where the roots were subjected to different treatments simultaneously (Figure 3E). We found that H_2_ could promoted the formation of proton-motive force across the membrane. The effects of H_2_ on ion fluxes that probably due to H_2_-uptake activities (i.e. H_2_→2H^+^ + 2e^−^). The electrochemical gradient can be converted into chemical energy for storage under the appropriate conditions. According to the acid growth hypothesis, elongation of plant cells can be promoted by H^+^ secretion and rapid acidification of the extension-limiting cell wall [30]. Therefore, the H_2_-uptaking activity of PM vesicles in *V. radiata* may provide a possible explanation for how H_2_ promotes hypocotyls elongation.

In addition to the effects on plant development, H_2_ also plays an important role in plant resistance to abiotic stress, such as salt tolerance [11]. The discovery of reducing Na^+^ influx (Figure 3J, 3K), may partly account for why H_2_ treatment attenuates salinity-induced growth inhibition in Arabidopsis [11].

### Hypothetical schematic diagrams of H_2_ metabolism in higher plants PM

According to our present knowledge, we propose a schematic diagram of H_2_ metabolism (H_2_-evolving and H_2_-uptaking) in PM vesicles of higher plants (Figure 4). Hydrogen acts directly as a proton and an electron donor. The flow of electrons and protons in carrying energy is the basis of all biosynthesis in life, which is inseparable from the coupling of oxidation and reduction reactions. The “valve” effect of bidirectional hydrogenases can play a very important role in regulating the energy homeostasis of plants. The discoveries in this study may provide some new insights about the molecular mechanisms of H_2_ and how to interpret its broad biological effects. The PM hydrogenase protein and its functions should be identified and analyzed in the future.

**Figure 4.**
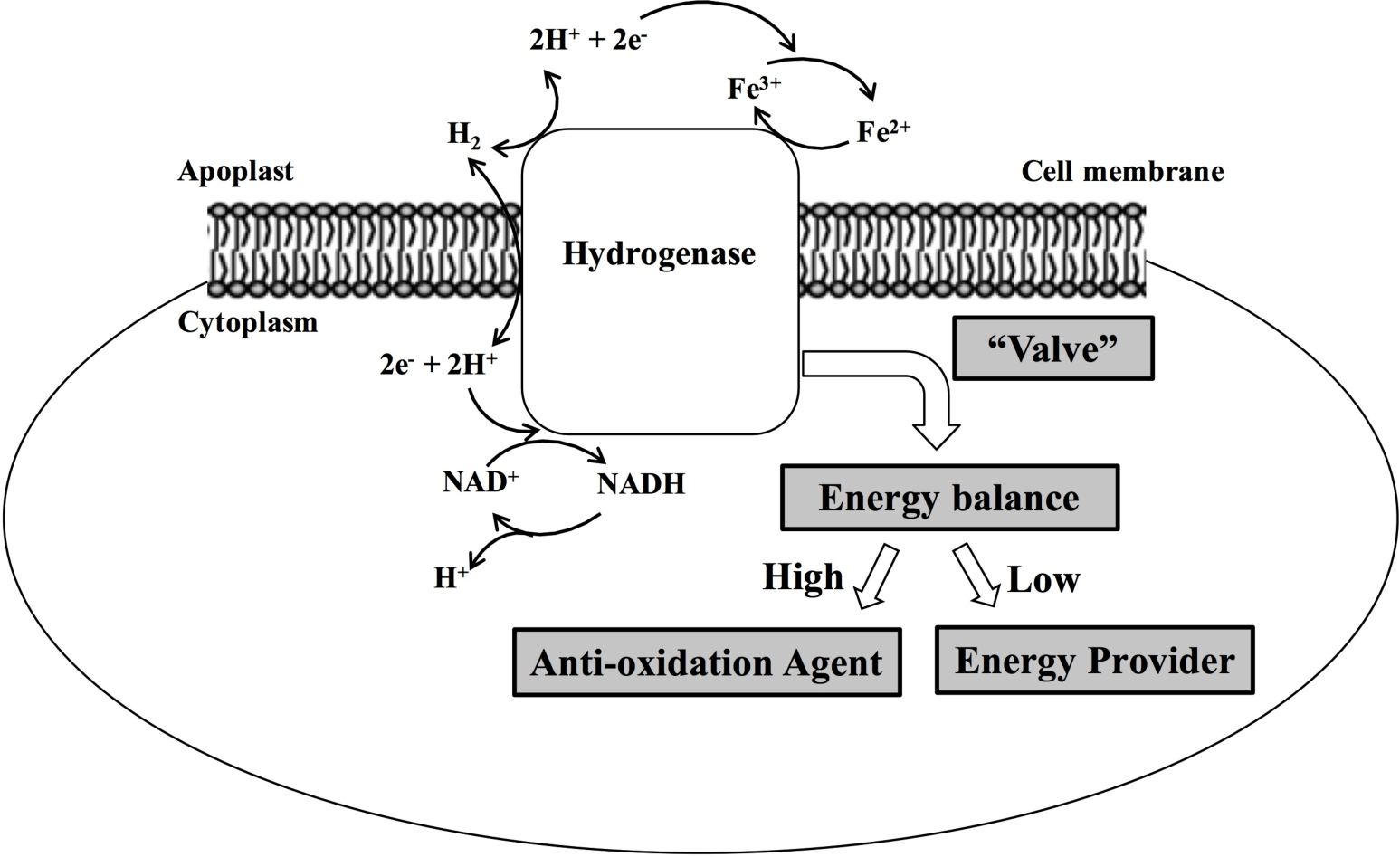
Hypothetical schematic diagrams of hydrogenase in higher plants PM

## Materials and methods

### Preparation of hydrogen-rich solutions

Purified H_2_ (99.999%) was generated by ZK200-hydrogen-producing apparatus from Zhongke Huiheng technology development Co. Ltd. (Beijing, China), then bubbled into ddH_2_O or required solution at a rate of 220 mL min^−1^ in 0.1 MPa for at least 20 min until H_2_ concentration reached saturation. Afterwards, the H_2_-saturated solution (0.8 mM) was immediately diluted to 0.2 mM, 0.4 mM or 0.68 mM. All H_2_-rich solution was freshly prepared. The H_2_ dissolved in solution was measured using a needle-type H_2_ sensor (Unisense, Aarhus N, Denmark).

### Plant materials

Mung beans (*Vigna radiata* [L.] R. Wilczek) were soaked in sterile ddH_2_O for 8 hours and grew in ddH_2_O at 23±1°C in darkness. Hypocotyls of 60-hour-old *V. radiata* were harvested for PM preparation. Intact plants or 5-mm long segments of hypocotyls (cut 5 mm below the cotyledons) were used for ion flux measurements. *C. annuum* were purchased from Shouguang Hongxiang Horticulture Co., Ltd. Stems of 4-week-old seedlings of *C. annuum* were harvested for PM preparation. To investigate the effects of hydrogen-rich water (HRW) on growth of *V. radiata*, the seeds of *V. radiata* were cultured in HRW containing 0.8 mM H_2_ for 3 days and continued to culture in H_2_-free water for another 2 days. Seedlings cultured in H_2_-free water were regarded as the negative control. Finally, the length and diameter of hypocotyls were measured. Assays were repeated with at least three independent experiments.

### Preparation of plasma membrane

The PM vesicles from *V. radiata* and *C. annuum* were prepared and purified as described previously with minor modifications [31]. The crude PM vesicles were dissolved in the suspended buffer containing 5 mM potassium phosphate buffer (pH 7.8), 5mM KCl, 0.25 M sucrose, 0.5% BSA(W/V) and 1 mM DTT. The PM vesicles were purified from microsomal fractions by partitioning in an aqueous polymer two-phase system. The ratio of hypocotyls weight to two-phase weight is 32: 8. The final pellet was re-suspended in the suspended buffer containing 10% (W/V) mannitol, put into liquid nitrogen immediately and stored at −80 °C until used. All the operations of extraction processes were performed at 4°C. Protein concentration was measured using Bradford protein assay kit (Tiangen Biotech, Beijing Co., LTD, CN). The purity of the PM was determined by standard marker assays[32]. Table S1 illustrated that the PM purity was enough to be used for further assay.

### Measurement of H_2_ production of the PM

H_2_ evolution of crude PM was monitored by headspace gas chromatography (GC) (Shimadzu, GCMS-QP2010S). A total volume of 500 µL reaction mixture in 1.9 mL gas chromatographic flasks containing about 2 mg crude PM and suspended buffer. The hypoxia conditions were obtained by flushing with nitrogen (N_2_) into the headspace of GC flasks for 2 minutes to remove O_2_. To study the effects of EDTA on H_2_ production, the reaction mixture was added with or without 1mM EDTA-Na_2_.

H_2_ produced by purify PM and dissolved in reaction solution was detected continually with a needle-type H_2_ sensor. H_2_ evolution was monitored in a total volume of 200 µL reaction mixtures. Potassium phosphate buffer (100 mM, pH 7.0) containing 2.5 mM MV was first bubbled with N_2_ to remove the O_2_, and then 80 µg purified PM vesicles (PM pretreated with 0.05% tritonX-100 and 5 µg NADH) were added. The reaction was initiated by addition of 10 mM Na_2_S_2_O_4_, which was freshly prepared with oxygen removed by sparging with N_2_. The PM vesicles boiled for 10 min served as the negative control.

### H_2_-dependent BV (benzyl viologen) reduction

H_2_ uptake hydrogenase activity (H2-dependent reduction of BV) was assayed spectrophotometrically at room temperature according to Pinske C at 578 nm [33] except that the buffer was 100 mM potassium phosphate (pH 7.0). H_2_ oxidation was recorded by the absorbance change of reduced BV. The final volume of 100 µL reaction mixture including 100 mM H_2_-saturated potassium phosphate (pH 7.0), 4 mM oxidized BV, and 10-20 µg PM (pre-treated with 0.05% tritonX-100 for 10 minutes on ice). Before reaction, either 5 µM NADH was added (to activate the PM), or nothing was added. The reaction was started by adding of 0.5 mM sodium dithionite (Na_2_S_2_O_4_). The PM vesicles that were boiled for 10 minutes served as the negative control. Assays were repeated with three independent experiments.

### NFORase enzymatic assay

NADH-ferricyanide oxidoreductase (NFORase) activity was tested at room temperature in the absence or presence of 0.2 mM or 0.68 mM H_2_. The activity was monitored by microplate readerVICTOR^3^ (Perkin Elmer Life Sciences, Waltham, MA, USA) as the time-dependent A420 and A340 absorbance change of ferricyanide (ε420 = 1.0 mM^−1^·cm^−1^) and NADH (ε340 = 6.22 mM^−1^·cm^−1^), respectively. Firstly, the reaction mixture containing 1.5 mM K_3_[Fe(CN)_6_] and reaction buffer (10 mM KCl, 3 mM MgSO4, 50 mM Tris-HCl, pH 7.5) were added into 96-well plates to detect absorbance. After approximately 5 minutes of measurement, the reaction mixture was added by premixed 0.2 mM NADH and 10-20 µg PM vesicles (pre-treated with or without 0.05% tritonX-100 on ice for 10 minutes) in the final volume of 100 µL. The group without the addition of PM was used as blank control. The NFORase activity was recorded over approximately a 45-min time course. Detailed conditions of assay are provided in the legends of the figures. Data are presented as standard deviation of the mean. Results of activity measurements were repeated twice in independent experiments.

### NADH regeneration

H_2_-driven reduction of NAD^+^ to NADH was detected spectrophotometrically at 340 nm [34]. NADH regeneration was monitored in a total volume of 100 µL of H_2_-saturated Tris-HCl buffer (50 mM, pH 8.0) containing 0.4 mM NAD^+^. Before adding to the reaction mixture, the PM vesicles of *V. radiata* hypocotyls were pre-treated with 0.05% tritonX-100 for 10 minutes on ice and activated by 5 µM NADH [18]. The reaction was started by adding 10-20 µg PM. PM was not added in the blank control. Results of activity measurements presented in this paper were repeated twice in independent experiments.

### Measurement of net ion flux with the NMT system

Net ion fluxes were measured noninvasively using Non-invasive Micro-test Technology (NMT; NMT100-SIM-YG, Younger USA LLC, Amherst, MA01002, USA). The gradients of ion concentration were measured by moving the ion-selective microelectrode between two positions close to the hypocotyl surface of approximately 8 mm below the cotyledons in a preset excursion (20 µm and 50 µm from the hypocotyl surface, along an axis perpendicular to the hypocotyls). Sixty-hour old *V. radiata* plant’s seedlings or excised hypocotyls were equilibrated in measuring solution (0.1 mM KCl, 0.1 mM CaCl_2_, 0.1 mM MgSO_4_, 0.1 mM NaCl, 0.3 mM MES, pH 6.0) for 10 min. Intact plants were placed in the 90 mm three-compartment plastic petri dishes. The hypocotyls and roots were placed in separate cells. A small notch was made in the barrier, which was between two cells to hold the seedlings (Figure 3E). The hypocotyls were soaked in fresh measuring solution and roots of intact plants were immersed in H_2_-free ddH_2_O or H_2_-free 150 mM NaCl solution. Stable net fluxes of H^+^, Na^+^, or K^+^ ions were measured for approximately 5 minutes. The H_2_-free ddH_2_O or NaCl was then replaced by ddH_2_O containing 0.8 mM H_2_ or NaCl containing 0.8 mM H_2_, respectively. The H_2_-free measuring solution impregnated with the excised hypocotyls was replaced by solution containing 0.2mM H_2_ or 0.8 mM H_2_. The net ion fluxes on the hypocotyls were measured for approximately another 5 minutes. Five to seven repetitions were carried out for the ion fluxes detection. All measurements of ion fluxes were carried out at Xuyue (Beijing) Sci. & Tech. Co., Ltd. (Beijing, China).

### Statistical Analysis

Unpaired t-test (two-tailed) was used to compare data from two groups. Data are expressed as mean ± SD. Differences with P-values <0.05 were considered significant. GraphPad Prism 6.0 was used for data analysis and figure creation.

## Compliance and ethics

The authors declare that they have no conflicts of interest with the contents of this article.

## Acknowledgments

We thank Fei Guo of the Institute of Pathogen Biology, Chinese Academy of Medical Sciences & Peking Union Medical College Research Center, for generous equipment support of PM extraction, and Jian Li, a member of their laboratory for helpful assistance. This work was supported by Special Fund for Agro-scientific Research in the Public Interest (201303023), Beijing Postdoctoral Research Foundation (2017-ZZ-019), Chaoyang District Postdoctoral Research Foundation (2018ZZ-01-11) and National Natural Science Foundation of China (81602408).

