## supplementary materials for "Hydrogen evolution and absorption phenomena in plasma membrane of higher plants"

**Table S1** Marker enzyme activities

| Marker enzyme | Inhibitor | Activity（μmol min$^{-1}$ mg $^{-1}$ pro.） |
|---|---|---|
| Plasma membrane ATPase | $Na_3VO_4$ | 0.376±0.024[a] |
| Tonoplast ATPase | $NaNO_3$ | 0.057±0.025[a] |
| Mitochondrial membrane ATPase | $NaN_3$ | 0 |
| Acid phosphatases | $Na_2MoO_4$ | 0 |

[a] indicated Mean±SEM

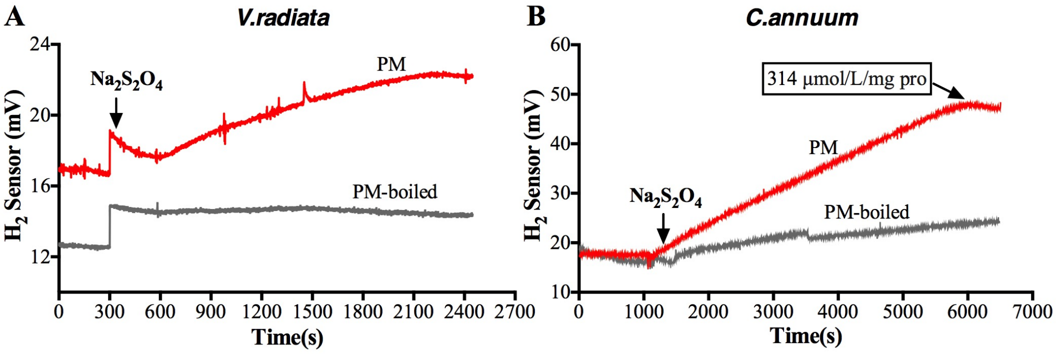


**Figure S1** $H_2$-evolution hydrogenase activities of purify PM vesicles were detected continuously by $H_2$-sensor. (A) $H_2$ evolution activity of purified PM vesicles from *V.radiata* hypocotyls. (B) $H_2$ evolution activity of purified PM vesicles from *C. annuum* stems.

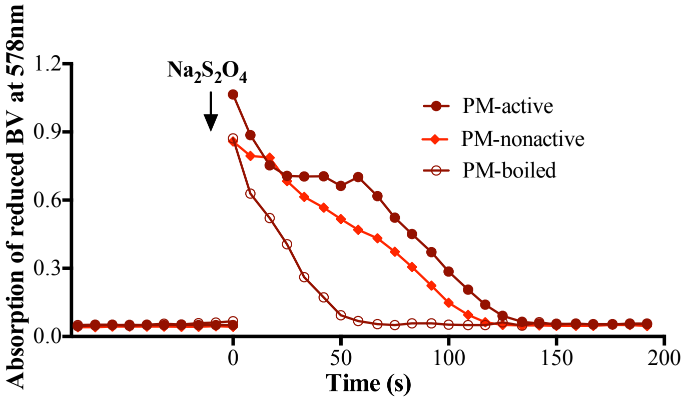


**Figure S2** The time-course of $H_2$ uptake activities of purified PM vesicles from *C. annuum* stems. PM-boiled, the purified PM boiled for 10 minutes served as the negative control. PM vesicles were either active or non-active by either adding or not adding 5 μM NADH. $Na_2S_2O_4$ initiated the reactions as indicated by arrows.

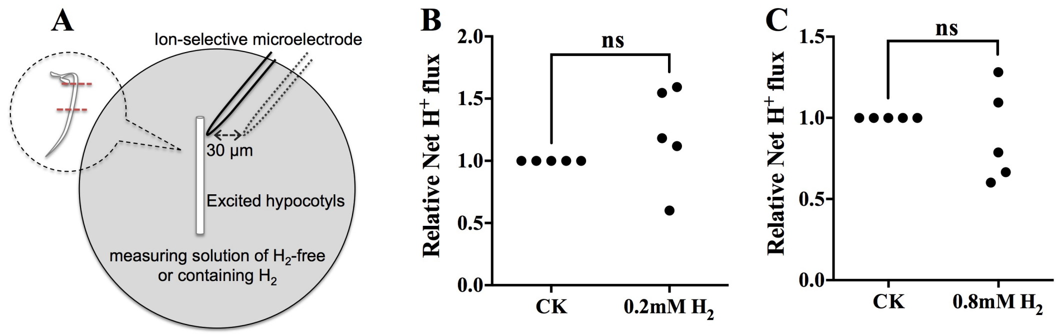


**Figure S3**. The effects of $H_2$ on net $H^+$ fluxes of excised *V.radiata* hypocotyls. (A) Schematic diagram of net $H^+$-fluxes measurement of hypocotyls from excised *V. radiata* seedlings by NMT. (B) Relative net ion fluxes of excised hypocotyls treated by 0.2mM $H_2$. (C) Relative net ion fluxes of excised hypocotyls treated by 0.8mM $H_2$. Each dot represented one sample.
